# RNA viruses drove adaptive introgressions between Neanderthals and modern humans

**DOI:** 10.1101/120477

**Authors:** David Enard, Dmitri A Petrov

## Abstract

Neanderthals and modern humans came in contact with each other and interbred at least twice in the past 100,000 years. Such contact and interbreeding likely led both to the transmission of viruses novel to either species and to the exchange of adaptive alleles that provided resistance against the same viruses. Here, we show that viruses were responsible for dozens of adaptive introgressions between Neanderthals and modern humans. We identify RNA viruses—specifically lentiviruses and orthomyxoviruses—as likely drivers of introgressions from Neanderthals to Europeans. Our results imply that many introgressions between Neanderthals and modern humans were adaptive, and that host genetic variation can be used to understand ancient viral epidemics, potentially providing important insights regarding current and future epidemics.

**One Sentence Summary:** Once out of Africa, modern humans inherited from Neanderthals dozens of genes already adapted against viruses present in their new environment.

## Main text

After their divergence 500,000 to 800,000 years ago, modern humans and Neanderthals interbred at least twice: the first time around 100,000 years ago (*1*) and the second around 50,000 years ago (*2*-*6*). The first interbreeding episode left introgressed segments of modern human ancestry within Neanderthal genomes, as revealed by sequencing of a single Altai individual (*7*). The second interbreeding episode left introgressed segments of European Neanderthal ancestry within the genomes of non-African modern humans. This history of interbreeding raises several important questions. Which introgressed sequences persisted in each species’ genome by chance and which persisted due to positive selection? And for those introgressed sequences that were positively selected, which pressures in the environment present at the time of interbreeding drove adaptation?

We recently found that viruses have driven as much as 30% of protein adaptation during evolution of the human lineage since divergence with chimpanzees, as well as a substantial fraction of adaptation in mammals overall (*8*). Because viruses were responsible for so much adaptation in humans, and because it is plausible that when Neanderthals and modern humans interbred they also transmitted viruses either directly or via their shared environment, we hypothesized that introgressions between Neanderthals and modern humans might have been driven by positive selection against the detrimental effects of viruses acquired from the other species. Specifically, we hypothesized a “poison/antidote” model of introgression whereby one species provides the other with both the “poison” in the form of a novel virus and the “antidote” in the form of virus-interacting alleles that confer resistance to these viruses. Consistent with our model, several cases of known adaptive introgressions from Neanderthals to modern humans involve specialized immune genes (*9*-*15*).

To test this model, we investigated the pattern of introgressions within a set of virus-interacting proteins (VIPs) (*8*). Adaptive introgressed segments are expected to be both longer and present in the population at higher frequencies than neutral introgressions (*16*). After interbreeding, adaptive Neanderthal haplotypes are expected to have rapidly increased in frequency in modern humans before recombination could break them down. Despite subsequent recombination, this should leave a distinct signature in modern human genomes. Specifically, we expect to observe regions of contiguous Neanderthal ancestry across multiple individuals that are longer and more frequent than expected under neutrality. Sankararaman et al. (*5*) identified segments of Neanderthal ancestry in the genomes of East Asian and European modern humans and estimated the frequencies of these segments. Therefore, we tested our hypothesis by examining whether VIPs are overrepresented among long, high-frequency introgressed segments relative to carefully matched non-VIPs (proteins not known to interact with viruses) (Materials and Methods). We analyzed East Asian and European modern human populations separately because they may have distinct histories of interbreeding with Neanderthals (*17*, *18*).

We focused on 4534 VIPs (~20% of the human proteome; Table S1) that engage in defined physical interactions with many viruses, including 20 human viruses with at least 10 VIPs (Table S2, Materials and Methods). VIPs were annotated based on interactions with modern viruses, but can be thought of as proxies for related viruses in ancient populations. This extension is supported by the fact that related viruses tend to use similar host VIPs (*8*). For example, VIPs interacting with HIV are also likely to interact with other lentiviruses. If enrichment of adaptive introgressions of HIV-interacting VIPs was observed, this could thus be taken as evidence of past adaptation related to a lentivirus rather than HIV itself.

Of the 4534 VIPs, 1920 VIPs were identified by low-throughput approaches and hand-curated from the virology literature (LT-VIPs), whereas 2614 VIPs were identified by high-throughput approaches (*19*) (HT-VIPs; Materials and Methods). Previously, we used a smaller set of 1256 LT-VIPs that are conserved across mammals to demonstrate that VIPs tend to be unusually highly conserved (*8*). Before studying patterns of introgression between modern humans and Neanderthals, we first demonstrated that our previous findings are consistent across this expanded set of VIPs. Using the full set of 4534 human VIPs, we show that compared to non-VIPs, VIPs exhibit (i) a lower average ratio of nonsynonymous to synonymous polymorphisms (Table S3); (ii) a higher proportion of rare, likely deleterious polymorphisms, reflected in the more negative values of Tajima’s D; and (iii) a higher density of functional and possibly deleterious segregating variants inferred by FUNSEQ (Table S3; Materials and Methods) (*20*-*23*). VIPs are also (i) found in regions of the genome with higher densities of coding sequences (*24*), regulatory sequences (*25*) and conserved genomic segments (*26*); (ii) are more highly expressed (*27*); and (iii) have more interacting protein partners in the network of human protein–protein interactions than non-VIPs (Table S3) (*28*, *29*). Note that while LT-VIPs and HT-VIPs have similar GC content, levels of evolutionary constraint, and densities of coding, regulatory, and conserved elements, they differ with respect to number of interacting partners, recombination rates (*30*, *31*), and abundance of FUNSEQ functional variants (Table S4). Thus, although we combined these two groups into a single VIP category for the rest of this analysis, we also systematically confirmed that the major results obtained when combining all VIPs also held true when using only the hand-curated LT-VIPs. In summary, we could confirm that the 4534 VIPs are more conserved and have more segregating deleterious variants than non-VIPs.

Next, in order to determine whether VIPs are enriched in segments introgressed between modern humans and Neanderthals, we first matched the regions containing VIPs and non-VIPs for levels of segregating deleterious variants. This was essential because we expected introgressed segments containing many deleterious polymorphisms to be preferentially eliminated by purifying natural selection, as demonstrated for introgressions both from Neanderthals to humans (*5*, *32*, *33*) and from humans to Neanderthals (*1*). To this end, we first defined all the genomic parameters that differed between introgressed and non-introgressed regions in both directions, including GC content, the number of human protein-protein interactions and multiple parameters controlling for levels of deleterious variants: Tajima’s D, FUNSEQ score, and densities of coding, regulatory, and conserved elements (Materials and Methods; Table S3 and Figs. S1, S2, S3 and S4). Because all of these genomic parameters also varied between VIPs and non-VIPs (Table S3), we used a bootstrap test to first match the VIPs with control non-VIPs for all relevant factors (Fig. S1 and Tables S5 and S6; Materials and Methods). We also systematically matched VIPs and non-VIPs with similar recombination rates in the bootstrap test (Materials and Methods) as adaptation in regions of low recombination should be hindered due to strong linkage between the adaptive variant and deleterious variants on the long introgressed haplotypes (*32*, *33*). We also measured the difference in proportion of adaptive introgressions at VIPs as a function of the recombination rate (*30*), as this difference should be more precisely quantified in regions of high recombination. Having determined how recombination and other genomic factors affect the occurrence of introgressions, we tested for an excess of introgressions from Neanderthals to modern humans.

We detected a strong and highly significant excess of Neanderthal introgressed segments encompassing VIPs in both East Asian and European populations. The enrichment becomes more pronounced for longer and more frequent introgressed segments both genome-wide and when restricting to regions of high recombination (Fig. 1A and B). These results are also robust when restricting analysis to LT-VIPs (Fig. S5). The excess of high-frequency introgressions was significantly greater in high-recombination regions of the genome (hypergeometric test using introgressed segments larger than 100 kb and at frequencies higher than 15%; East Asia *P*=0.016, Europe *P*=0.039) (Fig. 1B). Overall we identified 121 (vs. 66 expected) segments longer than 100 kb overlapping VIPs in East Asia (bootstrap test *P*<10^-3^) and 103 (vs. 68 expected) in Europe (*P*<10^-3^), implying that roughly 35-45% of all introgressed haplotypes containing VIPs have been subject to positive selection in modern humans. For the introgressions that are long (≥100 kb) and frequent (≥15%), the absolute counts are smaller but the enrichment is even more pronounced: 36 (vs. 11) segments in Asia (*P*<10^-3^), and 19 (vs. 6) in Europe (*P*<10^-3^), implying that ~70% of all high-frequency and long introgressions containing VIPs were adaptive. Altogether, these results demonstrate that adaptive introgression in response to viruses is far more prevalent than previously known based on the small number of examples (*9*-*15*).

**Figure 1.**
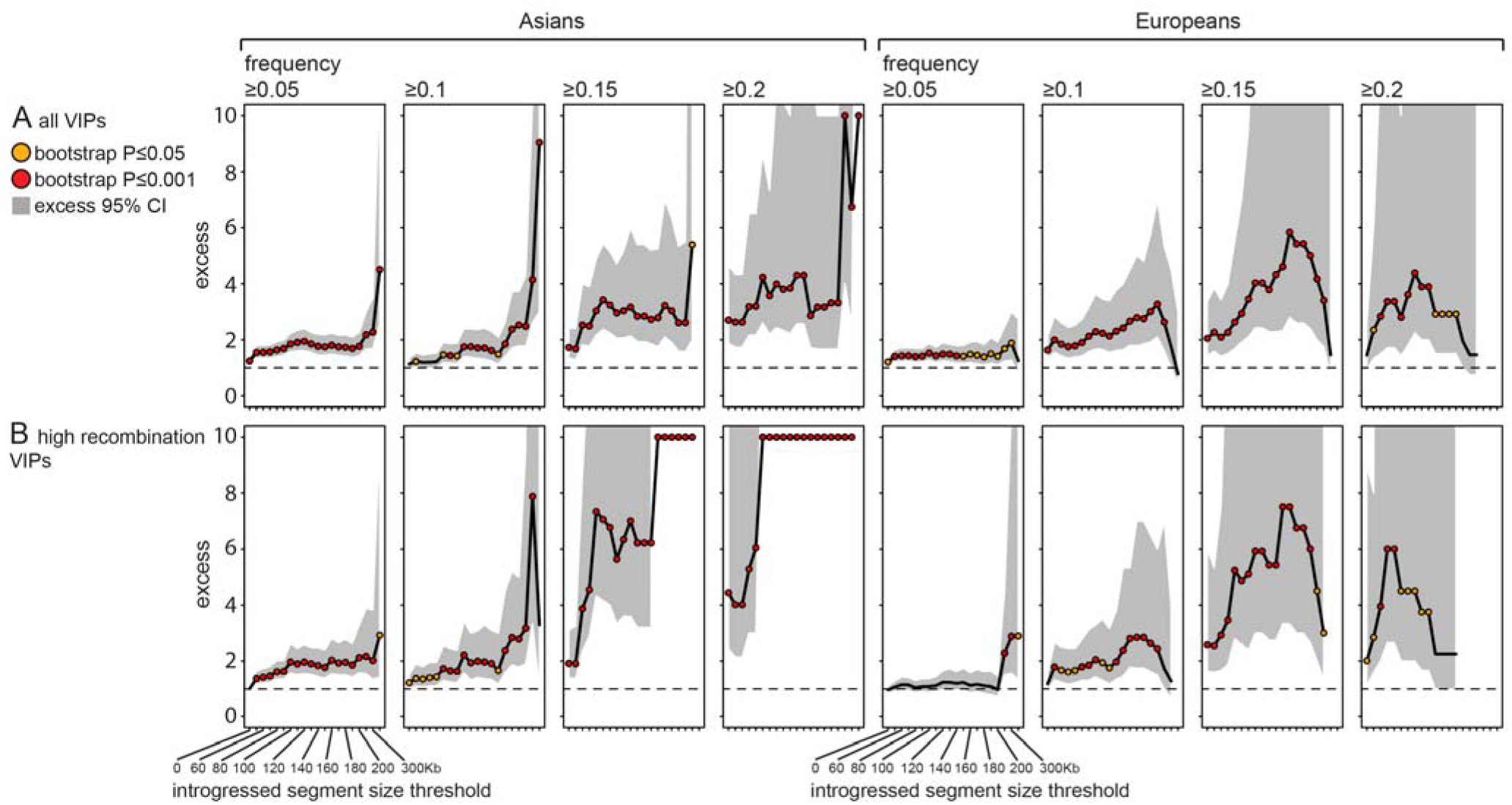
Excess of introgressions from Neanderthals to modern humans at VIPs. The graphs show the relative excess (y-axis) of introgressed segments of Neanderthal ancestry within Asian and European modern human genomes as a function of increasing lower segment size threshold (x-axis) and increasing lower segment frequency threshold (panels form left to right). An excess of 4 indicates that there are 4-fold more introgressed segments overlapping VIPs than there are introgressed segments overlapping matched control non-VIPs, according to the bootstrap test (Materials and Methods). The black line represents the observed excess, and the grey area represents the 95% confidence interval of the excess. For representation purposes, any excess greater than 10 is depicted as 10 in the graphs. Segment size thresholds for which the confidence interval is not represented correspond to thresholds beyond which there are no introgressed segments overlapping control non-VIPs. Orange dots: bootstrap test *P*<0.05. Red dots: bootstrap test *P*<0.001. The dashed line indicates an excess of 1, i.e., the case in which equal numbers of segments overlap VIPs and control non-VIPs. The lower segment size threshold was increased until there were fewer than three remaining introgressed segments overlapping VIPs or non-VIPs included in the matching. A) Excess in all VIPs across the entire genome. B) Excess in VIPs across high recombination regions of the genome (recombination rate > 1.5 cM/Mb, the median recombination rate within introgressed segments).

We next tested for an excess of introgressions from modern humans to Neanderthals, using the data on introgressed genomic regions into a single Altai Neanderthal individual from Kuhlwilm et al. (*1*). Although the introgressed segments in a single individual necessarily represent only a fraction of all introgressions, they should be biased towards more frequent introgressions, somewhat improving our statistical power with regard to adaptive introgressions. Because adaptive introgressed segments are expected to be larger than neutral ones, we estimated the excess of segments of modern human ancestry in the single Altai Neanderthal individual genome at VIPs as a function of their size (Fig. 2A and B). We found a large excess of long segments of modern human ancestry at VIPs (Fig. 2A). Furthermore, as predicted, the excess is more pronounced in high-recombination regions of the genome (Fig. 2B) (hypergeometric test *P*=0.045). We confirmed that this excess was also detected using only high-quality LT-VIPs (Fig. S6).

**Figure 2.**
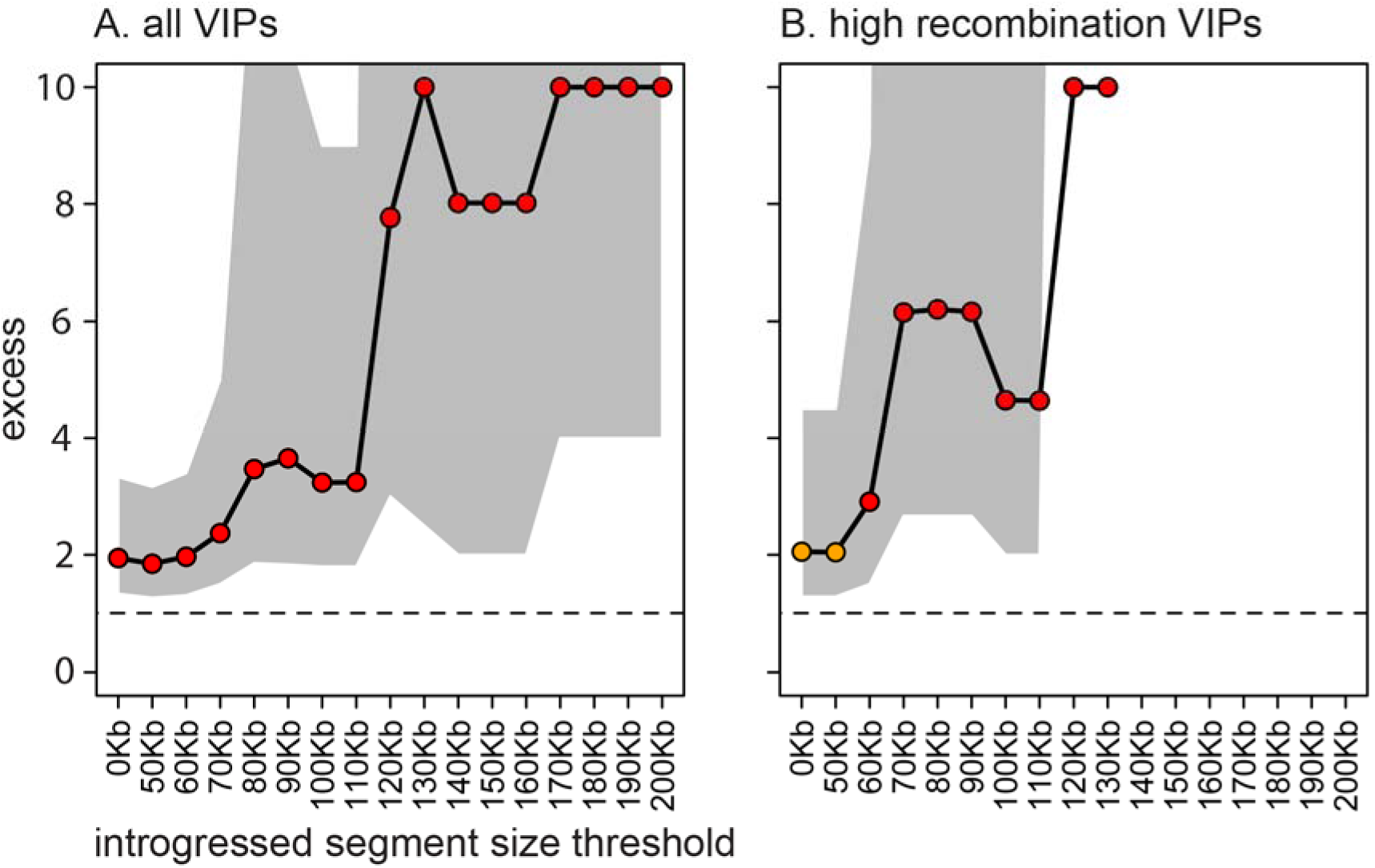
Excess of introgressions from modern humans to Neanderthals at VIPs. Legend as in Fig. 1. A) All VIPs. B) High recombination VIPs.

We next asked whether it is possible to identify ancient viruses that drove the adaptive introgressions from Neanderthals to modern humans by testing whether VIPs associated with specific viruses were more likely to be introgressed overall and also be present in long and frequent introgressions. While such an analysis in the direction from modern humans to Neanderthals is severely underpowered with only 19 VIPs found in introgressed segments over 100 kb in the Altai Neanderthal, the number is much larger in modern humans with 152 VIPs found in long (≥100 kb) and frequent (≥15%) Neanderthal-introgressed haplotypes, thus providing increased statistical power to test in this direction.

The 20 modern human viruses that interact with ten or more VIPs are used as proxies for the ancient related viruses that infected humans at the time of interbreeding (Table S2). We found no significant heterogeneity in the representation of these 20 viruses among introgressions overall either in Europeans (permutation test *P*=0.42; Materials and Methods) or in East Asians (*P*=0.12). We did, however, detect significant heterogeneity specifically in long (≥100kb) and high-frequency (≥15%) introgressions in Europeans (*P*=0.009), but not in East Asians (*P*=0.1).

We then asked whether ancient RNA or DNA viruses were more likely to have been involved, with some expectation that because RNA viruses are more likely to jump from one species to another (*34*, *35*), they should also be more likely to drive adaptive introgressions. The 20 viruses well-represented in our VIP set are almost evenly distributed between RNA viruses (2684 VIPs) and DNA viruses (2547 VIPs) (Table S2). Of the 2684 RNA VIPs, 1563 interact with only RNA viruses, while out of 2547 DNA VIPs, 1426 interact with only DNA viruses. A total of 1121 VIPs interact with both RNA and DNA viruses. In order to determine whether introgressions were skewed towards either RNA or DNA viruses, we used the bootstrap test to compare the number of introgressions at VIPs that interact with only one RNA virus with the number of introgressions at VIPs that interact with only one DNA virus and are located far from any RNA VIP (≥500 kb; Supplementary Text).

Consistent with the lack of overall heterogeneity among the 20 well-represented viruses, we did not detect any significant skew in East Asia (Fig. 3A). By contrast, in Europe we detected a remarkable bias of long, high-frequency introgressions towards RNA VIPs (Fig. 3A) compared to DNA VIPs. This pattern was much more pronounced for VIPs with high recombination rates (Fig. 3B). In European modern humans, we found 16 high-frequency (≥15%) introgressed segments containing VIPs that interact with a single RNA virus in high-recombination regions of the genome, versus only 1.6 expected by chance (bootstrap test *P*<10^-3^; Methods). These results were further confirmed using only LT-VIPs (Fig. S7A and B).

**Figure 3.**
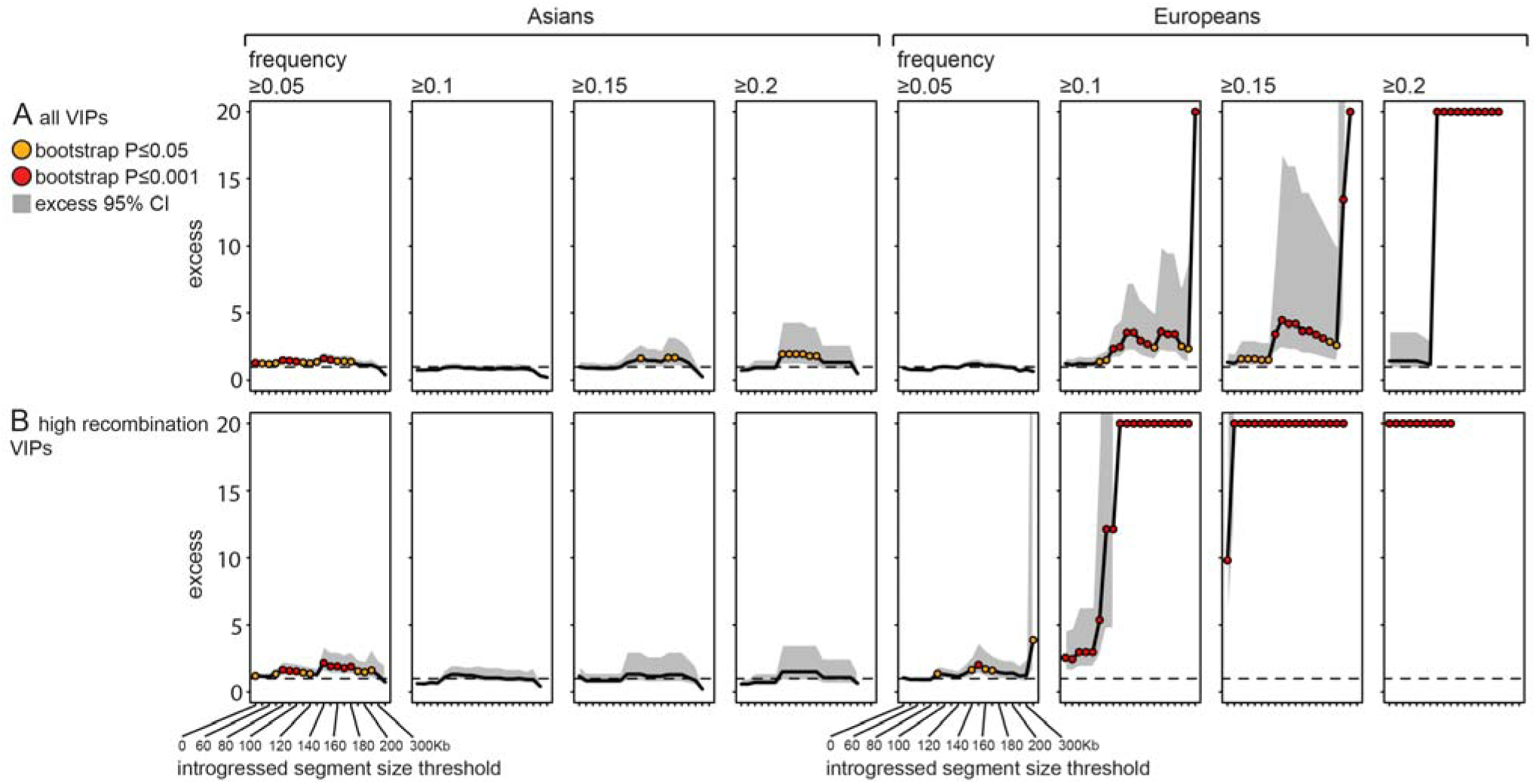
Excess of introgressions from Neanderthals to modern humans at RNA VIPs vs. DNA VIPs. Legend as in Fig. 1, except that the y-axis represents the excess of introgressions at RNA VIPs vs. DNA VIPs, rather than VIPs vs. non-VIPs.

We next tried to identify which families of ancient RNA viruses explain the observed skew towards RNA VIPs in Europeans. To this end, we used only VIPs specific for a single RNA virus. Of the 11 RNA viruses included in this analysis (Table S2), only HIV, Influenza A virus (IAV) and Hepatitis C virus (HCV) overlap with more than five introgressed segments in Europeans. Therefore, only ancient lentiviruses, orthomyxoviruses, or flaviviruses related to HIV, IAV or HCV, respectively, could possibly explain the skew towards RNA VIPs. It appears that both HIV-only and IAV–only VIPs were each responsible for a large excess of high-frequency, long adaptive introgressed segments in European modern humans compared to individual DNA viruses (Fig. S8A, B, C and D). As expected, the excess was particularly strong for HIV-only and IAV-only VIPs within high-recombination regions (Fig. S8B and D). Within high-recombination regions, we found seven (vs. 0.29 expected by chance) high-frequency (≥15%) introgressed segments overlapping IAV-only VIPs (*P*< 10^-3^) and eight (vs. 0.83 expected) overlapping HIV-only VIPs (P<10^-3^). Table S7 lists the specific VIPs found in these introgressed segments. These results were robust when restricting to HIV-only LT-VIPs (Fig. S7C and D). However, the small number of IAV-only LT-VIPs (56 overall vs. 195 for HIV, and only 15 in high recombination regions) did not provide sufficient power to detect a significant excess of introgressions, as indicated by subsampling of HIV-only LT-VIPs to the number of IAV-only LT-VIPs (bootstrap test *P*>0.05 for IAV-only LT VIPs and all ten random subsamples and all introgression lengths and frequencies).

Although we detected no significant enrichment of introgressed segments at HCV-only VIPs (Fig. S8E and F), this might be an issue of power since HCV has far fewer unique VIPs than HIV and IAV (157 versus 405 and 490, respectively). Subsampling of HIV-only and IAV-only VIPs to the number of HCV-only VIPs results in low power to detect a significant excess (Fig. S9), leaving the possibility open that HCV-like viruses might also have been involved. Taken together, these results suggest that European modern humans suffered from infections by at least two or possibly more ancient RNA viruses. More specifically, they suggest that European modern humans suffered from infections by ancient lentiviruses and orthomyxoviruses related to HIV and IAV, respectively, and using similar VIPs as HIV and IAV at the time of contact with Neanderthals. The introgressions of VIPs from Neanderthals likely provided a measure of protection against these viruses. The resulting burst of adaptive introgressions is still detectable in European modern humans.

Here, we presented evidence that introgressions from Neanderthals to modern humans and vice versa are strongly enriched for proteins interacting with viruses. We also showed that this enrichment was more pronounced for long and frequent introgressions in regions of high recombination. In the case of introgressions from Neanderthals to modern humans, we detected a particularly strong signal for VIPs interacting with several RNA viruses in Europeans, including HIV and IAV. Overall these results provide support for the “poison-antidote” hypothesis under which the interactions between modern humans and Neanderthals exposed each species to novel viruses while gene flow between the species afforded a measure of resistance, by allowing VIPs that were already adapted to the presence of specific viruses to cross the species boundaries.

This study opens many avenues for future investigation. For instance, the fact that introgressed VIP alleles (“antidotes”) from one human species were adaptive in the other implies previous encounters of the donor species with the specific viruses (“poisons”) in their past. However, it remains to be determined whether these viruses were transmitted directly from Neanderthals to modern humans and vice versa, or from other animals present in the environment. Sequencing of additional Neanderthal genomes might help us identify specific viruses transferred to Neanderthals through contact with modern humans. It is also possible that other pathogens in addition to viruses might have played a role in adaptive introgressions between these two species. Finally, it should be extremely fruitful to study whether the presence of Neanderthal ancestry at VIPs still leads to variable susceptibility to modern viruses in modern humans.

Our results also suggest the possibility that adaptation was an integral part of the interbreeding between Neanderthals and modern humans. Indeed, based on the measured excess of introgressions at VIPs, we estimate that of all long (≥100 kb) and high-frequency (≥15%) introgressed segments from Neanderthals to modern humans, 32% (54 of 171) in Asians and 25% (27 of 105) in Europeans were positively selected in response to viruses (Methods). Given that adaptation due to resistance to viruses is only a part of all possible sources of adaptation, and that in fact other cases of adaptive introgressions likely unrelated to viral resistance have been found (*36*), we thus conclude that a substantial proportion of all introgressions was likely adaptive.

Our results imply that the genomes of humans and plausibly many other species contain abundant signals of numerous past arms races with diverse viruses and other pathogens, raising the possibility of using host genomic sequence data to study ancient epidemics. The results of such studies could provide important insights into the dynamics of past, present, and future epidemics.

## Acknowledgments

We wish to thank Rajiv McCoy, Jamie Blundell, Kerry Geiler-Samerotte, Sandeep Venkataram, Emily Ebel and other current and former members of the Petrov Lab for comments on the manuscript. This work was supported by the grants NIH 1RO1GM10036601 and NIH R35GM118165 to D.A.P. The authors declare no competing interest.

## Supplementary Materials

Materials and Methods

Supplementary Text

Table S1-S7

Fig S1-S9

